# Decreasing Serine Levels During Growth Transition Triggers Biofilm Formation in *Bacillus subtilis*

**DOI:** 10.1101/564526

**Authors:** Jennifer Greenwich, Alicyn Reverdy, Kevin Gozzi, Grace Di Cecco, Tommy Tashjian, Veronica Godoy-Carter, Yunrong Chai

**Author notes:** These two authors contributed equally to this work (Jennifer Greenwich, Alicyn Reverdy).

## Abstract

Biofilm development in *Bacillus subtilis* is regulated at multiple levels. While a number of known signals that trigger biofilm formation do so through the activation of one or more sensory histidine kinases, it was recently discovered that biofilm activation is also coordinated by sensing intracellular metabolic signals, including serine starvation. Serine starvation causes ribosomes to pause on specific serine codons, leading to a decrease in the translation rate of *sinR*, which encodes a master repressor for biofilm matrix genes, and ultimately biofilm induction. How serine levels change in different growth stages, how *B. subtilis* regulates intracellular serine levels in response to metabolic status, and how serine starvation triggers ribosomes to pause on selective serine codons remain unknown. Here we show that serine levels decrease as cells enter stationary phase and that unlike most other amino acid biosynthesis genes, expression of serine biosynthesis genes decreases upon the transition into stationary phase. Deletion of the gene for a serine deaminase responsible for converting serine to pyruvate led to a delay in biofilm formation, further supporting the idea that serine levels are a critical intracellular signal for biofilm activation. Finally, we show that levels of all five serine tRNA isoacceptors are decreased in stationary phase compared to exponential phase. Interestingly, the three isoacceptors recognizing UCN serine codons are reduced to a much greater extent than the two that recognize AGC and AGU serine codons. Our findings provide evidence for a link between serine homeostasis and biofilm development in *B. subtilis*.

**IMPORTANCE:** In *Bacillus subtilis*, biofilm formation is triggered in response to various environmental and cellular signals. It was previously proposed that serine limitation acts as a proxy for nutrient status and triggers biofilm formation at the onset of biofilm entry through a novel signaling mechanism caused by global ribosome pausing on selective serine codons. In this study, we revealed that serine levels decrease at the biofilm entry due to catabolite control and a shunt mechanism. We also show that levels of five serine tRNA isoacceptors are differentially decreased in stationary phase compared to exponential phase; three isoacceptors recognizing UCN serine codons are reduced much greater than the two recognizing AGC and AGU codons. This indicates a possible mechanism for selective ribosome pausing.

## INTRODUCTION

Bacteria in the natural environment are often found in surface-attached multicellular communities known as biofilms (1–3). Biofilms are a leading cause of hospital-acquired infections (4). Biofilms also pose problems in the environment and industrial settings. *Bacillus subtilis* is a Gram-positive, rod-shaped bacterium commonly used as a model system for studies of biofilm formation (5, 6). Under laboratory settings, *B. subtilis* is capable of forming two different types of biofilms: pellicles, which form at the air-liquid interface, and colony biofilms, which form on solid surfaces (2, 5). *B. subtilis* is also able to form plant root-associated biofilms when living in the rhizosphere (7–9). The genetic circuitry governing biofilm formation in *B. subtilis* has been well-studied (reviewed in (6)), but less is known about the signals and signal transduction mechanisms that trigger biofilm formation.

Canonically, extracellular signals activate one or more sensory histidine kinases, which then triggers a phosphorelay, leading to the phosphorylation of Spo0A, a master regulator for cell development in *B. subtilis* (10–12)(Fig. 1A). Upon phosphorylation, Spo0A~P activates transcription of the *sinI* gene (13). *sinI* encodes an antagonist for the master biofilm repressor SinR (14, 15). SinI binds to SinR and sequesters it off its DNA targets, leading to expression of SinR-repressed genes responsible for production of the biofilm matrix, which include a 15-gene *epsA-O* and a three-gene *tapA-sipW-tasA* operons (Fig. 1A)(6). Interestingly, small changes in SinR protein levels were shown to have dramatic effects on the expression of matrix genes (13, 16), primarily due to the ultra-sensitivity of the SinI-SinR regulatory module, indicating that its levels need to be tightly regulated within the cell. Recently, metabolic stimuli of biofilm formation have been identified (16–18). In a previous study, we showed that serine starvation acts as a proxy for nutrient limitation at the onset of stationary phase and triggers biofilm formation in *B. subtilis* (16). Serine starvation results in decreased translation of *sinR* due to ribosomes preferentially pausing on the four UCN (N stands for A, G, U, C) serine codons, and that the four UCN codons are overly abundant in the *sinR* gene (16). It remains to be confirmed if serine levels decrease during the transition from exponential to stationary phase growth. This also leads to other interesting questions such as how *B. subtilis* cells regulate serine levels and how decreased serine levels cause ribosomes to pause on selective serine codons.

**Figure 1:**
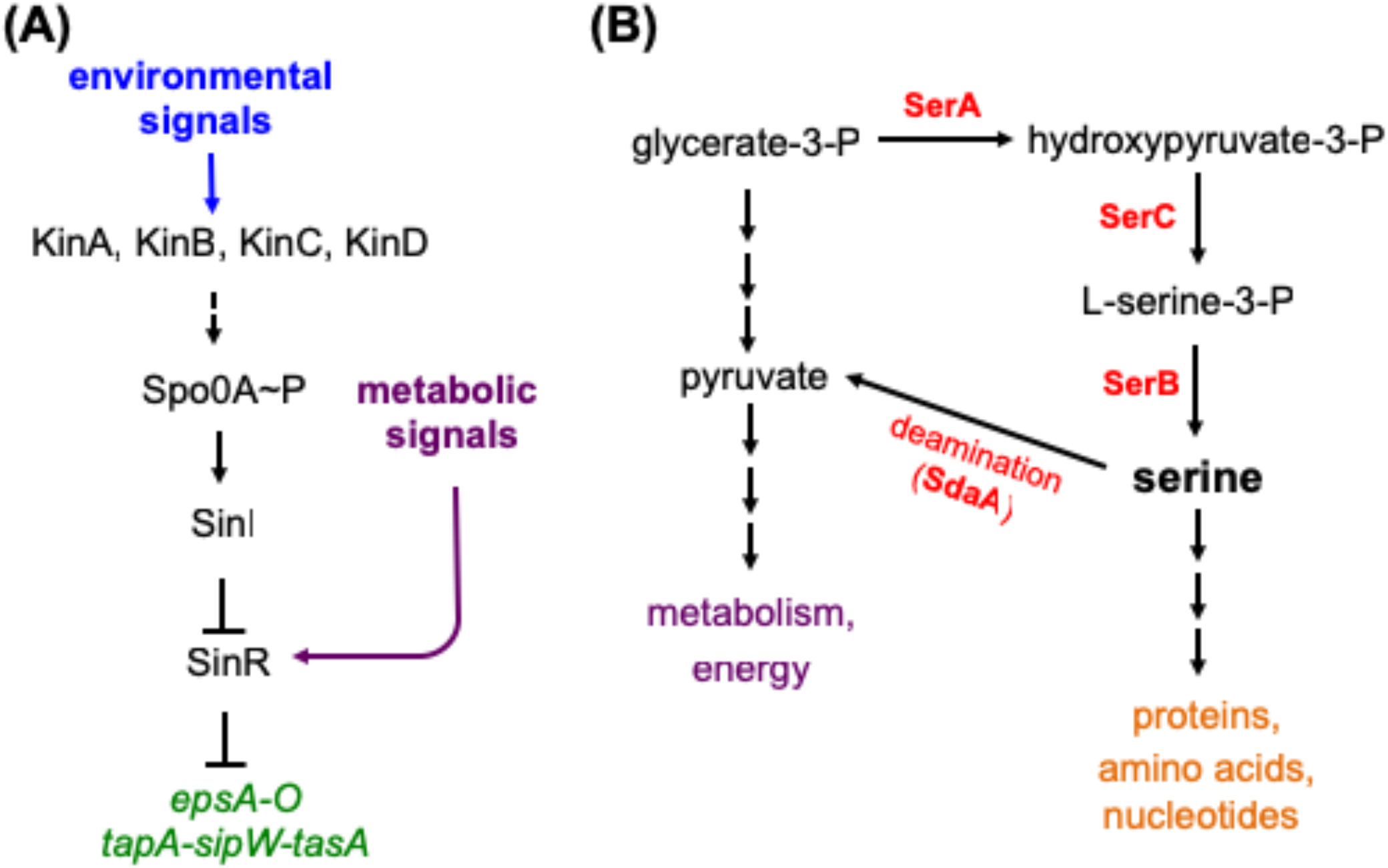
The genetic circuitries governing biofilm formation and serine metabolism in *Bacillus subtilis*. **(A)** The genetic circuitry of biofilm formation in *B. subtilis* has been elucidated. There are multiple layers of regulation, which allow cells to fine-tune biofilm formation prior to commitment. Extracellular signals activate one or more histidine kinases, leading to a phosphorelay, which phosphorylates Spo0A. Upon phosphorylation, Spo0A~P is active and in turn activates transcription of *sinI*, which encodes an antagonist for the master biofilm regulator, SinR. SinI antagonizes SinR repression thus allowing expression of the biofilm operons *epsA-O* and *tapA-sipW-tasA*. **(B)** Serine metabolism is linked to central metabolism through glycerate-3-phosphate (G-3-P). G-3-P is a precursor for serine biosynthesis through the activities of SerA, SerC, and SerB. SerA carries the rate-limiting step in serine biosynthesis. The SdaA deaminase complex is involved in converting serine back to G-3-P. While the pathway is known, the regulation of the genes involved remains elusive.

The amino acid serine is a centrally important biomolecule, not only as a building block for protein synthesis, but also as a precursor of nucleotides, other amino acids (such as cysteine, tryptophan, and glycine), and phospholipids (19). While the biochemical reactions involved in serine metabolism are well known, how bacterial cells regulate serine homeostasis and maintain intracellular serine concentrations remains poorly understood. Serine biosynthesis is intimately linked to central metabolism (Fig. 1B)(20). Glycerate-3-phosphate serves as both a glycolytic intermediate and a precursor to serine (20). The first, rate-limiting step in serine biosynthesis is the conversion of glycerate-3-phosphate to 3-phosphohydroxypyruvate, catalyzed by SerA (Fig. 1B)(20). The regulation of *serA* transcription has been elucidated in *Escherichia coli*, and interestingly enough, *serA* is regulated by global metabolic regulators including CRP (<u>c</u>AMP <u>R</u>eceptor <u>P</u>rotein) and Lrp (<u>L</u>eucine <u>R</u>ich Repeat <u>P</u>rotein), both of which activate *serA* transcription (21). CRP, also known as CAP (<u>C</u>atabolite <u>A</u>ctivator <u>P</u>rotein), activates genes responsible for secondary carbon source utilization in *E. coli* in a cAMP-dependent fashion (22, 23). Lrp responds to the amino acid leucine in the cell and is activated under leucine starvation (24). This indicates that under nutrient starvation, transcription of *serA* is activated in *E. coli*. In *B. subtilis*, regulation of *serA* expression on a genetic level remains unclear. Different from *E. coli*, in *B. subtilis*, carbon metabolism is chiefly regulated in a cAMP-independent manner by CcpA (<u>C</u>atabolite <u>C</u>ontrol <u>P</u>rotein <u>A</u>) (25, 26). A link between carbon metabolism and biofilm formation has been investigated in several recent studies (27, 28). Biofilm formation is triggered when cells are starved for glucose, which is dependent on catabolite control, as deletion of the *ccpA* gene increases biofilm formation (27). This indicates that CcpA negatively regulates biofilm formation in *B. subtilis* (27). Another global metabolic regulator CodY was also shown to be involved in biofilm formation in *B. subtilis* (29). In both cases, the underlying mechanisms remain unknown.

In bacteria, serine levels are also affected by a single enzymatic reaction converting serine to pyruvate and ammonia (Fig. 1B)(30). This reaction is carried out by a serine deaminase (also known as serine dehydratase) (30, 31). Serine deamination is clearly important because in *E. coli*, there are three serine deaminases and at least one of the three serine deaminases is active under any given growth condition (32). In *B. subtilis*, only one serine deaminase has been identified (33). This two-subunit protein complex is encoded by the *sdaAA* and *sdaAB* genes (33). The metabolic benefits(s) of serine deamination have not been fully investigated and remain largely unclear. Previous studies have shown that serine is depleted far faster than other amino acids in *E. coli* upon nutrient limitation or when cells enter stationary phase (34–36). Presumably, serine is converted to pyruvate and used for central metabolism, energy generation, and even gluconeogenesis. This can be considered as a shunt pathway for serine from being used as a building block for protein synthesis. It is unclear how *B. subtilis* cells would regulate serine homeostasis to ensure an adequate balance between carbon flow into amino acid biosynthesis and central metabolism.

In this study, we set out to elucidate the mechanism by which serine depletion triggers biofilm formation and how cells regulate intracellular serine levels in *B. subtilis*. We found that serine levels are decreased in stationary phase relative to exponential phase, and that serine biosynthesis declines as cells enter stationary phase. The key serine biosynthetic gene *serA* is regulated in an indirect manner by catabolite control in *B. subtilis*, but unlike in *E. coli*, *serA* in *B. subtilis* appears to be expressed when nutrients are plentiful. In addition to a decrease in serine levels, the abundance of serine tRNAs (a total of 5 isoacceptors) that recognize six synonymous serine codons (UCA, UCC, UCG, UCU, AGC, and AGU) is also decreased in stationary phase compared to the exponential phase, but the decrease differs significantly for different isoacceptors.

## MATERIALS AND METHODS

### Strains, media, and growth conditions

For general purposes, *Bacillus subtilis* strains PY79, NCBI3610 (hereafter as 3610), and their derivatives were grown at 37°C in lysogenic broth (LB) (10 g tryptone, 5 g yeast extract and 10 g NaCl per liter broth) or on solid LB medium supplemented with 1.5% agar. For assays of biofilm formation, LBGM or MSgg media were used. LBGM is composed of LB with supplementation of 1% glycerol (v/v) and 100 μM MnSO_4_ (37). The MSgg medium is as previously described (5). *Escherichia coli* DH5α was used as a host for molecular cloning and was grown at 37°C in LB medium. When required, antibiotics were added at the following concentrations for growth of *B. subtilis*: 1 μg/ml of erythromycin, 5 μg/ml of tetracycline, 5 μg/ml of chloramphenicol, 100 μg/ml of spectinomycin, and 10 μg/ml of kanamycin. For growth of *E. coli*: 100 μg/ml of ampicillin. A list of strains used in this study is summarized in Table 1.

**Table 1:**
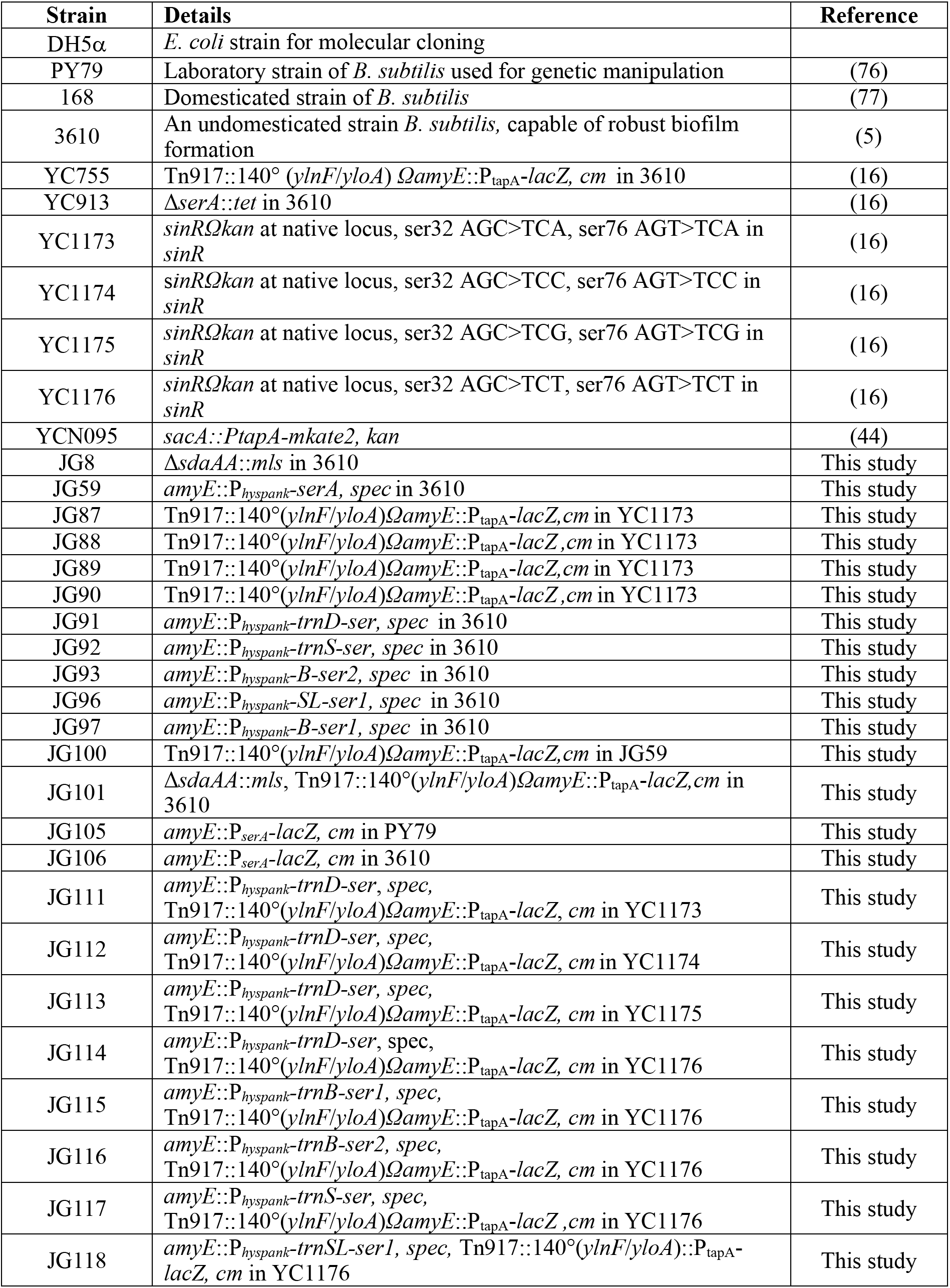
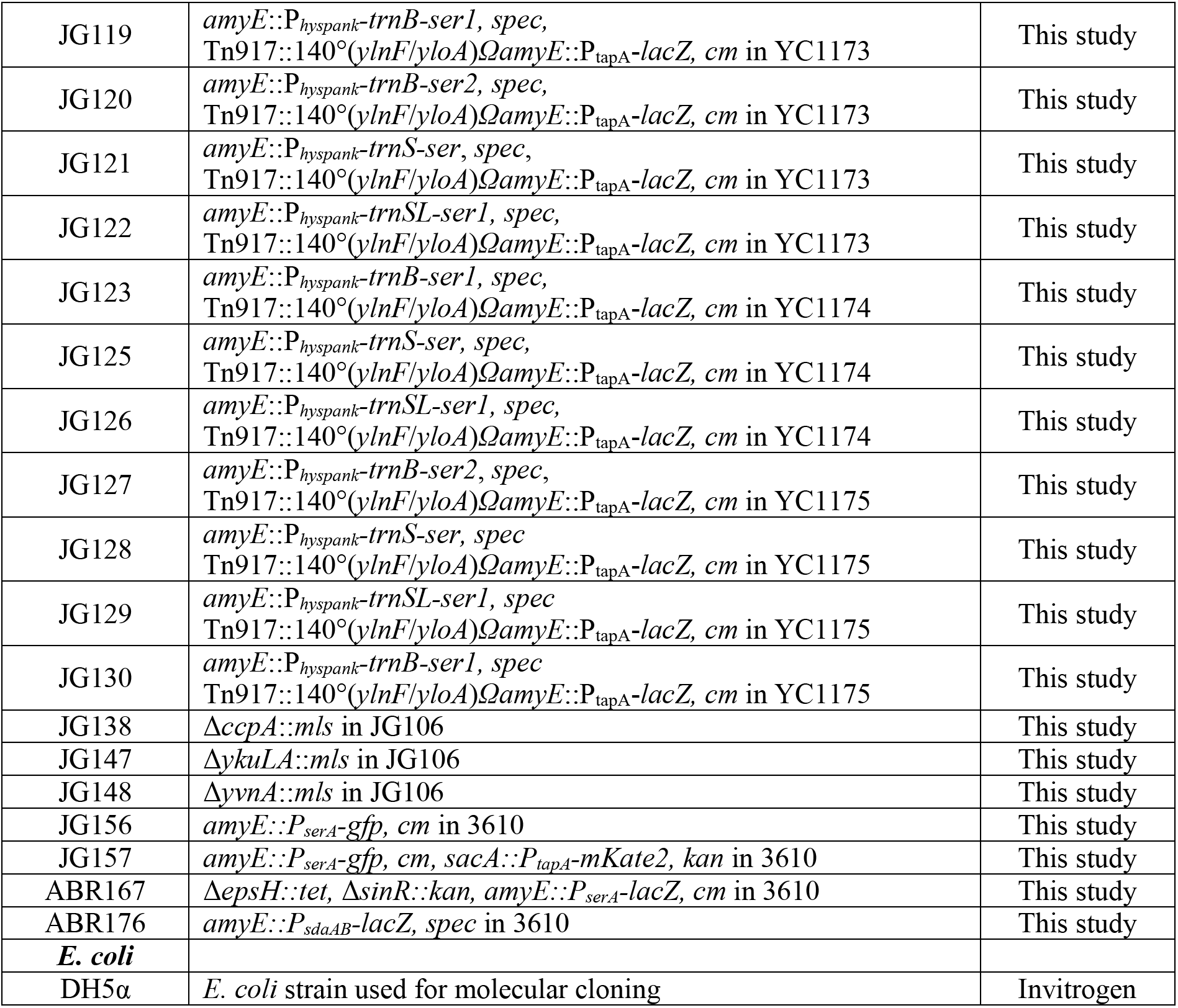
Strains Used in This Study

### DNA manipulation

General methods for molecular cloning followed the published protocols (38). Restriction enzymes (New England Biolabs) were used according to the manufacturer’s instructions. Transformation of plasmid DNA into *B. subtilis* strains was performed as described previously (39). SPP1 phage-mediated general transduction was also used to transfer antibiotic-marked DNA fragments among different strains (40). Plasmids used in this study are listed in Tables 2 and oligonucleotides (purchased from Eurofins) are listed in Table S1.

**Table 2:** Plasmids Used in This Study.

| Plasmid | Details | Reference |
| --- | --- | --- |
| pDG268 | An *amyE* integration vector with a promoter-less *lacZ*, amp, cm | BGSC |
| pDG1728 | An *amyE* integration vector with a promoter-less *lacZ*, amp, spec | BGSC |
| pDR111 | An *amyE* integration vector that contains *P<sub>hyspank</sub>*, amp, spec | BGSC |
| pYC121 | An *amyE* integration vector with a promoter-less *gfp-mut2*, amp, spec | (43) |
| pJG01 | *amyE::P<sub>hyspank-serA</sub>* in pDR111, amp, spec | This study |
| pJG02 | *amyE::P<sub>serA-lacZ</sub>* in pDG268, amp, cm | This study |
| pJG03 | *amyE::P<sub>hyspank-trnB-ser1</sub>* in pDR111, amp, spec | This study |
| pJG04 | *amyE::P<sub>hyspank-trnB-ser2</sub>* in pDR111, amp, spec | This study |
| pJG05 | *amyE::P<sub>hyspank-trnS-ser</sub>* in pDR111, amp, spec | This study |
| pJG06 | *amyE::P<sub>hyspank-trnSL-ser1</sub>* in pDR111, amp, spec | This study |
| pJG07 | *amyE::P<sub>hyspank-trnD-ser</sub>* in pDR111, amp, spec | This study |
| pJG08 | *amyE::P<sub>serA-gfp</sub>, amp, cm* | This study |
| pABR174 | *amyE::P<sub>sdaAB-lacZ</sub>* in pDG1728, amp, spec | This study |

### Strain construction

For construction of the insertional deletion mutations of *ccpA* and *sdaAA* in 3610, the corresponding mutants in the *B. subtilis* strain 168 were obtained from the Bacillus Genetic Stock Center (BGSC, www.bgsc.org) at Ohio State University. Antibiotic cassette marked deletions were introduced into 3610 using SPP1 phage-mediated transduction to generate strains JG138 and JG8, respectively (40).

To overexpress *serA* in *B. subtilis*, the *serA* coding sequence was amplified by PCR using 3610 genomic DNA as the template and primers serA-F and serA-R. The PCR product was then cloned into the *HindIII* and *BamHI* sites of pDR111, which contains an isopropyl β-D-1-thiogalactopyranoside (IPTG)-inducible *hyperspank* promoter flanked by the *amyE* gene generating the recombinant plasmid pJG01(41). The recombinant plasmid was then introduced into the *B. subtilis* laboratory strain PY79 by genetic transformation for a double-crossover recombination of the DNA sequences at the *amyE* locus. The construct was then introduced into 3610 by transformation to generate strain JG59 (40). Overexpression of each of the five tRNA^ser^ isoacceptor genes was done in a similar manner, using primers listed in Table S1 to produce strains JG91 through JG97.

To compare expression of the *serA* gene in the wildtype and various deletion strains, the regulatory promoter sequence of *serA* was amplified by PCR using primers PserA-F and PserA-R and genomic DNA of *B. subtilis* strain 3610 as the template. The PCR products were digested with EcoRI and *HindIII*, and cloned into the plasmid pDG268 (42), which carries a chloramphenicol-resistance marker and a polylinker upstream of the *lacZ* gene between two arms of the *amyE* gene, to make a P*_SerA_-lacZ* transcriptional fusion, resulting in the recombinant plasmid pJG02. For comparison of *tapA* expression in wild-type and various engineered strains for serine metabolism, lysate from a strain containing a P*_tapA-lacZ_* fusion (YC755) was introduced into strains JG08 and JG59 by SPP1 phage mediated transduction. The P*_tapA-lacZ_* fusion was introduced into a locus between the *ylnF* and *yloA* genes, at 140° on the chromosome, as previously described (13). This insertion keeps the *amyE* gene intact, which allows for additional insertions such as the P*_hyspank_-serA* or tRNA overexpression constructs, and for measurement of *lacZ* transcription as well as IPTG induction.

To examine single-cell fluorescence and track both *serA* expression and biofilm matrix gene expression, a dual reporter strain was constructed. The regulatory sequence of *serA* was amplified by PCR using primers PserA-F and PserA-R, and genomic DNA of *B. subtilis* strain 3610 as the template. The PCR product was digested with EcoRI and HindIII, and cloned into the plasmid pYC121, which has a promoter-less *gfp* gene *(gfp-mut2)* flanked by the *amyE* sequence (43). This resulting plasmid, pJG08, was then introduced into the *B. subtilis* laboratory strain PY79 by genetic transformation for a double-crossover recombination of the DNA sequences at the *amyE* locus. The reporter fusion was then moved into 3610 by transduction (40), generating strain JG156. To construct the dual-reporter, the reporter was introduced to YCN095, which contains a P*_tapA_-mKate2* reporter fusion integrated at *sacA* (44), generating strain JG157.

To measure *sdaAA* expression, a 150-bp promoter region (P*_sdaAB_*) was amplified using primers PsdaAB-F and PsdaAB-R and 3610 genomic DNA as the template. The PCR product was digested with BamHI and *HindIII* restriction enzymes and inserted onto the pDG1728 vector backbone to generate plasmid pABR174. pABR174 was then transformed into *B. subtilis* PY79 and further into 3610 to generate strain ABR176 (40).

### Serine auxotroph assay

To harvest supernatant, wild-type 3610 cells, or its derivatives were grown overnight on an LB plate at 37°C. Single colonies were grown overnight in 3 mL of LB medium under shaking conditions at 37°C. Cells were sub-cultured at 1:100 into MSgg medium. Supernatant was collected at OD_600_=0.6 (exponential phase) and OD_600_=1.4 (stationary phase), respectively. Supernatant was filter-sterilized, passed through a mini-sizing column (MW cutoff 5K, Qiagen), and then diluted in MSgg to normalize cell count to 1×10^8^ CFU/mL. To test auxotroph growth, the Δ*serA* strain (YC913) was grown overnight on an LB plate at 37°C. Single colonies were grown overnight in 3 mL of LB medium under shaking conditions at 37°C. Cells were sub-cultured at 1:100 into MSgg media for six hours. Cells were washed in fresh MSgg and then diluted to a concentration of approximately 100 CFU/mL in fresh MSgg. One mL of cells was added to 1 mL of supernatant. Cells were grown for 24 hours under shaking conditions at 37°C and plated. After 24 hours of incubation, cells were plated and CFU was counted the next day. Resulting CFUs were divided by starting CFU levels to assess relative growth ability. Experiments were performed in triplicate and repeated at least three times for each strain.

For generation of the standard curve, YC913 was grown as described above, however after 6hrs of growth, the culture was diluted to 10^3^. This dilution was inoculated 1:100 into 2mL culture tubes containing MSgg with varying concentrations of serine (Sigma). An initial inoculum was plated for CFU onto LB plates. The remaining cultures grew at 37°C under shaking conditions for 24hrs and were then plated for CFU. The calculation of the curves was standardized to the initial inoculum. The standard curve was carried out in triplicate for four independent experiments.

### β-Galactosidase activity assay

For assays of β–galactosidase activity, an established protocol was used (ref). Briefly, cells were incubated in a flask of MSgg or LBGM medium at 37°C with shaking. One milliliter of culture was collected at each indicated time point after inoculation. Cells were spun down and pellets were resuspended in 1 ml Z buffer (40 mM NaH_2_PO_4_, 60 mM Na2HPO4, 1 mM MgSO_4_, 10 mM KCl and 38 mM β–mercaptoethanol) supplemented with 200 μg/ml freshly made lysozyme. Resuspensions were incubated at 30°C for 15 min. Reactions were started by adding 200 μL of 4 mg/ml ONPG (2-nitrophenyl β–D-galactoside) and stopped by adding 500 μL of 1 M Na_2_CO_3_. Samples were spun down to remove any cell debris. OD_420_ and OD_550_ values of the samples were recorded using a BioRad SmartSpec 3000. The β-galactosidase-specific activity was calculated according to the equation: Activity (Miller Units) = (OD_420_-1.75(OD_550_)/(time × volume × OD_600_)) × 1000. Experiments were performed in triplicate and repeated at least three times.

### Pellicle biofilm and β-galactosidase activity assay

*B. subtilis* cells were grown in LB broth at 37°C to mid-log phase. For pellicle formation, 4 μL of the cells mixed with 4 mL of liquid MSgg media in 12-well plates (Corning). Plates were incubated at 30°C for up to 48 hours. Images were taken using a Leica MZ10F dissecting scope. For β-galactosidase activity assays on pellicle biofilm samples, pellicle biofilms were set up as described above and collected at 24 or 48 hours. Pellicles were resuspended using the media in the well and gently sonicated to disrupt cell chains and bundles, but not individual cells. One milliliter of the pellicle was spun down and resuspended in 1 mL Z buffer. 100 μL of sample was then diluted into 900 μL Z buffer due to high cell density in the pellicle. The β-galactosidase activity was assayed as described above. The activity was calculated similarly as described above. Experiments were performed in triplicate and repeated at least three times.

### Quantitative real-time PCR

Five hundred microliter (500 μL) of *B. subtilis* cells were harvested during exponential phase (OD_600_~0.6) and stationary phase (OD_600_~1.4), respectively, and added to one milliliter of RNAProtect Bacteria Reagent (Qiagen). After incubation at room temperature for 60 minutes, samples were spun down (15,000 g for 5 min) and the cell pellet was resuspended in one milliliter of Trizol Reagent (Thermo) and incubated for five minutes at room temperature. Total RNA was extracted using the Zymo Direct-zol RNA kit (Zymo) following the manufacturer’s protocol. RNA concentration and purity was measured using a NanoDrop One (Thermo). RNA was converted to cDNA using a high capacity cDNA reverse transcription kit (Applied Biosciences). qPCR was performed using the primers listed in table S3 using the protocol provided by the manufacturer for the Fast SYBR green master mix (Applied Biosystems) using a StepOnePlus real-time PCR machine (Applied Biosystems). Relative expression was calculated using the comparative threshold cycle (ΔΔCT) for each sample. Technical triplicates were run for each sample to calculate the ΔΔCT and biological triplicates were run for each strain, and the experiment was repeated three times, for a total of nine biological replicates. The relative expression was standardized using the endogenous 16S control with the exponential phase samples used as the reference sample. A one-way ANOVA was used to assess statistical significance (P<0.05).

### Dual reporter assay

For the dual reporter assay, 50 mL of MSgg was inoculated 1:100 from an overnight LB broth culture with strain JG157 containing the P*_serA-gfp_* and P*_tapA-mKate2_* fluorescent reporters. Cultures were grown in shaking at 37°C and cells were harvested at mid-exponential (~OD_600_=0.5), early stationary (~OD_600_=1.5), mid-stationary (OD_600_=3.0), and late stationary phases (after overnight growth). To visualize cells under the microscope, 1 mL of cells were harvested, washed with 1 mL of PBS buffer to remove residual LB medium, and then concentrated in 100 μL of PBS buffer. A volume of 2 μL of cells was added to a 1% agarose pad and covered with a coverslip. Cells were imaged using a Leica DFC3000 G camera on a Leica AF6000 microscope. Images of samples collected from different timepoints were taken using the same exposure settings. For GFP observation, the setting of the excitation wavelength was 450–490 nm, while the setting of the emission wavelength was 500–550 nm. For mKate2 observation, the excitation wavelength setting was at 540–580 nm and the emission wavelength setting was at 610–680 nm. The experiment was conducted in triplicate and images taken were representatives of the three replicates.

## RESULTS

### Serine Levels Decrease Upon Entry into Stationary Phase

We previously showed that early stationary-phase conditions (biofilm entry) or induced serine starvation in the exponential phase led to ribosome pausing on selective serine codons and decreased translation of *sinR* (16). We thus proposed that serine starvation occurs in early stationary phase and acts as a trigger for biofilm induction. It remained to be confirmed if serine levels were decreasing upon cells entering stationary phase under normal conditions. We initially applied biochemical approaches (primarily HPLC and MS/Spectrometry-based) to measure concentrations of free intracellular amino acids, but experienced inconsistency of the results in both experimental and control samples (data not shown). We then decided to develop a novel microbiological assay for the measurement (see Methods). Briefly, wild type *B. subtilis* cells capable of synthesizing serine were grown in a minimal medium (MSgg) without exogenous serine. Supernatant from either the exponential (OD_600_=0.6) or early stationary (OD_600_=1.4) phase culture was prepared as described in the Methods. Supernatants were then added to a fresh minimal medium that does not contain any exogenous serine and inoculated with a *B. subtilis* serine auxotroph strain (Δ*serA*)(16). Δ*serA* cells were cultured for 24 hours and then plated (Fig. 2A). Relative growth of the auxotroph was calculated by comparing the final colony forming units (CFU) to the initial inoculum. We also engineered two *B. subtilis* strains predicted to accumulate higher levels of serine (JG08 for *serA* overexpression or *serA*^++^, and JG59 for Δ*sdaAA*) and collected supernatants from those cultures accordingly. As shown in Fig. 2B, supernatant from exponentially growing wild type cells was able to support more than four times higher growth of the auxotroph strain than the stationary phase supernatant (as calculated by CFU, Fig. 2B). Interestingly, the exponential phase supernatant of JG08 (*serA*^++^) was able to support about ten times higher growth of the auxotroph than the supernatant from the stationary phase culture (*P*<0.0001, Fig. 2B), while the exponential phase supernatant from the Δ*sdaAA* strain (JG59) also demonstrated a significant increase in supporting auxotroph growth compared to supernatant from the stationary phase (*P*<0.05, Fig. 2B).

**Figure 2:**
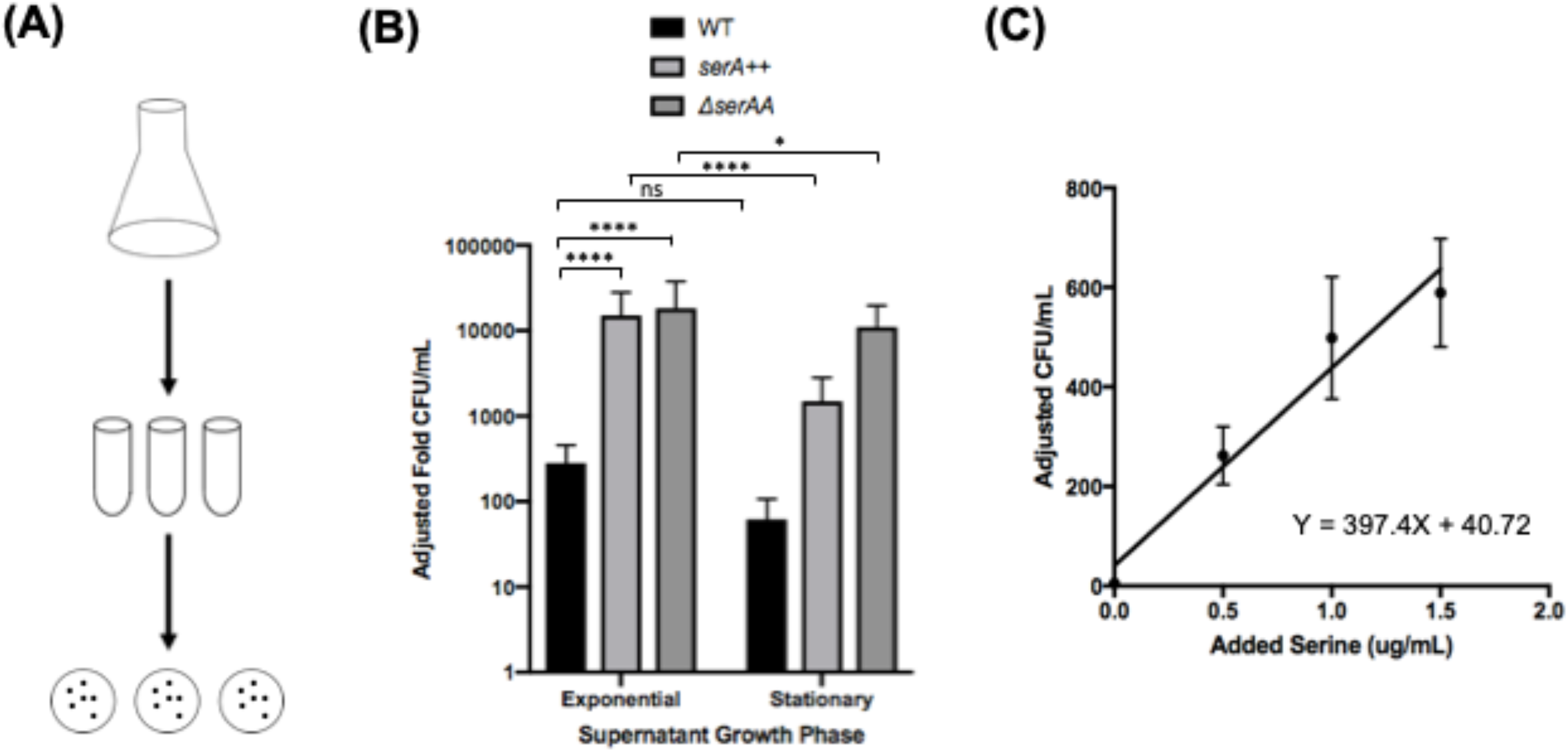
Supernatants of *B. subtilis* cells support growth of a serine auxotroph. **(A)** A schematic of the microbiological assay used to assess serine levels in the supernatant. Supernatant from strains capable of serine biosynthesis grown in MSgg medium was prepared during exponential phase (OD_600_=~0.6) and early stationary phase (OD_600_=~1.4) to mimic biofilm entry. Auxotroph cultures, *ΔserA* (YC913), were washed and sub-cultured for growth into the supernatants and then plated for CFU counts. Fold growth is relative to CFU of initial inoculum. **(B)** Exponential phase supernatant from cells with *serA* overexpression, *serA*+ +(JG08), was able to support growth about ten times better than stationary phase supernatant (****= p≤0.0001). Adding supernatant from also significantly increased population growth during exponential phase compared to stationary phase in from cells with a deleted *sdaAA* gene (JG59) (*= p≤0.05). Supernatant did not show a significant decrease in cell growth from wildtype supernatant in stationary compared to exponential stage (ns, p>0.05). Exponential phase supernatants of both JG08 and JG59 supported much higher growth of the auxotroph than the exponential phase supernatant of the wild type (****=p≤0.0001), indicating the effectiveness of genetic manipulation in increasing cellular serine accumulation. **(C)** Standard curve of Δ*serA*(YC913) in MSgg growth medium supplemented with different concentrations of serine. CFU count was adjusted to the CFU of initial inoculum. The best-fit curve has equation y= 397.40X+40.72 (R=0.9626) and can be used to better estimate the concentration of serine in the harvested supernatant.

When compared to auxotroph growth in the media with known concentrations of serine (Fig. 2C), we estimated about 0.82 μg/ml and 0.08 μg/ml serine in the exponential and early stationary phase of the wild type supernatants, 38.0 μg/mL and 3.65 μg/mL serine in the exponential and early stationary phase of JG08 (*serA*^++^) supernatants, and 80.99 μg/mL and 27.85 μg/mL serine in the exponential and early stationary phase of JG59 (Δ*sdaAA*) supernatants, respectively. Our results supported the note of significantly higher levels of free serine in the supernatant of the exponential phase culture than the stationary phase culture. Although we could not directly measure the intracellular serine levels, we infer that intracellular serine levels are likely proportionally higher in the exponential phase cells than in the stationary culture. Finally, when a saturating amount of exogenous serine (200 μM) was added to the media together with either the exponential or stationary phase supernatant, the auxotroph strain no longer demonstrated difference in growth, indicating that the reduced growth support by the stationary phase supernatant was not due to the presence of a killing factor (data not shown).

### Manipulation of Intracellular Serine Levels Alters Biofilm Formation

We previously showed that adding exogenous serine to the media delayed biofilm formation by *B. subtilis* (16). This suggested that manipulation of serine levels may affect the timing of biofilm induction. In an effort to understand the physiological implications of changes in intracellular serine levels during growth transition, we decided to investigate how manipulation of the serine biosynthesis or shunt pathway would affect biofilm formation. As described above, two enzymatic pathways play key roles in serine homeostasis in *B. subtilis*: *de novo* serine biosynthesis primarily from pyruvate mediated by SerA/SerB/SerC and deamination of serine back to pyruvate mediated by the serine deaminase complex SdaAA/SdaAB (Fig. 1B). Based on the microbiological assays described above (Fig. 2B), we already observed that supernatants from the two engineered strains (JG08 and JG59) supported substantially higher auxotroph cell growth compared to that of the wild type (*P*<0.0001 for both JG08 and JG59, respectively), indicating that both genetic manipulations successfully allowed increased serine accumulation. Our results thus suggest that overexpression of *serA* alone is able to generate much higher serine levels in the cell. In the case of Δ*sdaAA*, presumably this mutant strain was unable to convert serine into pyruvate, which likely led to increased intracellular serine levels.

We next investigated how those gene mutations may affect biofilm formation. We performed biofilm assays for the engineered strains and observed that deletion of the *sdaAA* gene severely delayed biofilm formation; the mutant formed a very weak pellicle biofilm at 24 hours while at 48 hours, the pellicle biofilm of the mutant showed up to a level comparable to the wild type (Fig. 3A). *sdaAA* encodes the alpha subunit of the two-protein deaminase complex. It should be noted that deletion of the beta subunit (SdaAB) showed a similar phenotype (data not shown), indicating that it is the complex responsible for this phenotype and not a moonlighting function of SdaAA. By using a β-galactosidase reporter assay, we further determined that transcription of the biofilm matrix gene *tapA* was decreased in the Δ*sdaAA* strain (Fig. 3B). At both 24 and 48 hours, there was a significant decrease in the expression of *tapA* relative to the wild type even though visible morphological difference in pellicle formation disappeared at 48 hours (Fig. 3A). On the other hand, overexpression of *serA* only mildly impacted biofilm formation with a modest delay at 24 hours (Fig. 3A) and changes in *tapA* expression by *serA* overexpression were also insignificant (Fig. 3B). It is possible that the increase in serine levels by *serA* overexpression is still below a threshold level to significantly alter biofilm formation. In the previous assay, we showed that the stationary phase supernatants from both *serA* overexpression and *sdaAA* supported much more autotroph growth compared to the WT, but to a lesser degree in the case of *serA* overexpression (Fig. 2B). Finally, overexpression of *serA* in the Δ*sdaAA* strain showed a very similar biofilm phenotype to that of the Δ*sdaAA* strain, indicating that serine deamination plays a more prominent role in serine homeostasis especially during growth transition (data not shown). We did not test how *ΔserA* impacted biofilm formation simply because the growth of the auxotroph strain required supplementation of large amounts of exogenous serine.

**Figure 3:**
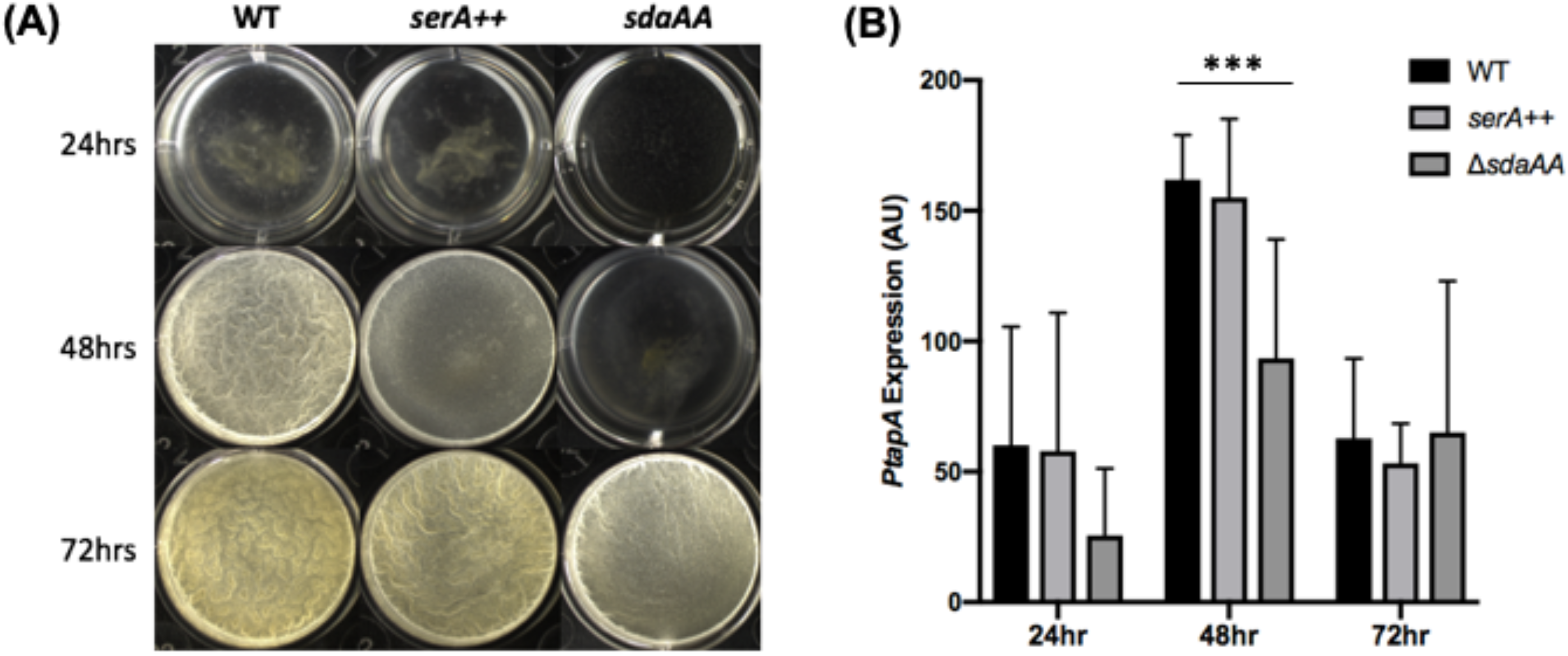
Inability to convert serine into pyruvate delays biofilm formation. **(A)** Δ*sdaAA*(JG08) shows a weaker biofilm robustness compared to wild type (3610) at 48 hours, but not at 72 hours in MSgg media, indicating a delay in biofilm formation. **(B)** *P_tapA_-lacZ* expression from pellicles grown in MSgg is significantly lower in Δ*sdaAA (*JG101) at 24 and 48 hours indicating a 24hr delay in biofilm production in this strain compared to wildtype (YC755) (***=p<0.001).

### *serA* Expression Decreases Upon Entry into Stationary Phase

We next wondered if decreasing serine levels upon cells entering stationary phase was due to the regulation of serine homeostasis. Amino acid biosynthesis genes are normally upregulated upon entry into stationary phase, as that is when cells resume *de novo* amino acid biosynthesis (45). This is the case for *serA* in *E. coli*, which is under catabolite control (21). To investigate if the expression pattern of *serA* is the same in *B. subtilis* as in *E. coli*, we constructed a P*_SerA_-lacZ* reporter fusion and tested *PserA* expression over time. As shown in Figs. 4A and 5A, in wild type cells, *serA* transcription initially increased until peaked at around three/four hours after inoculation depending on initial inoculum. Following this, *serA* transcription began to decrease. The peak of *serA* expression corresponded with the timing when cells began to transition from exponential phase to early stationary phase (OD_600_≈1.0). The timing also corresponded to when the induction of biofilm genes occurs based on previous studies (43, 46). A decrease in *serA* expression could contribute to the decrease in serine levels. More importantly, it indicates that in *B. subtilis*, serine biosynthesis is regulated in an unusual manner relative to biosynthesis of other amino acids. Decrease in *serA* expression upon entry into stationary phase in *B. subtilis* was also confirmed by RNA-seq in a recent study [(47) and DeLoughery A, personal communications].

**Figure 4:**
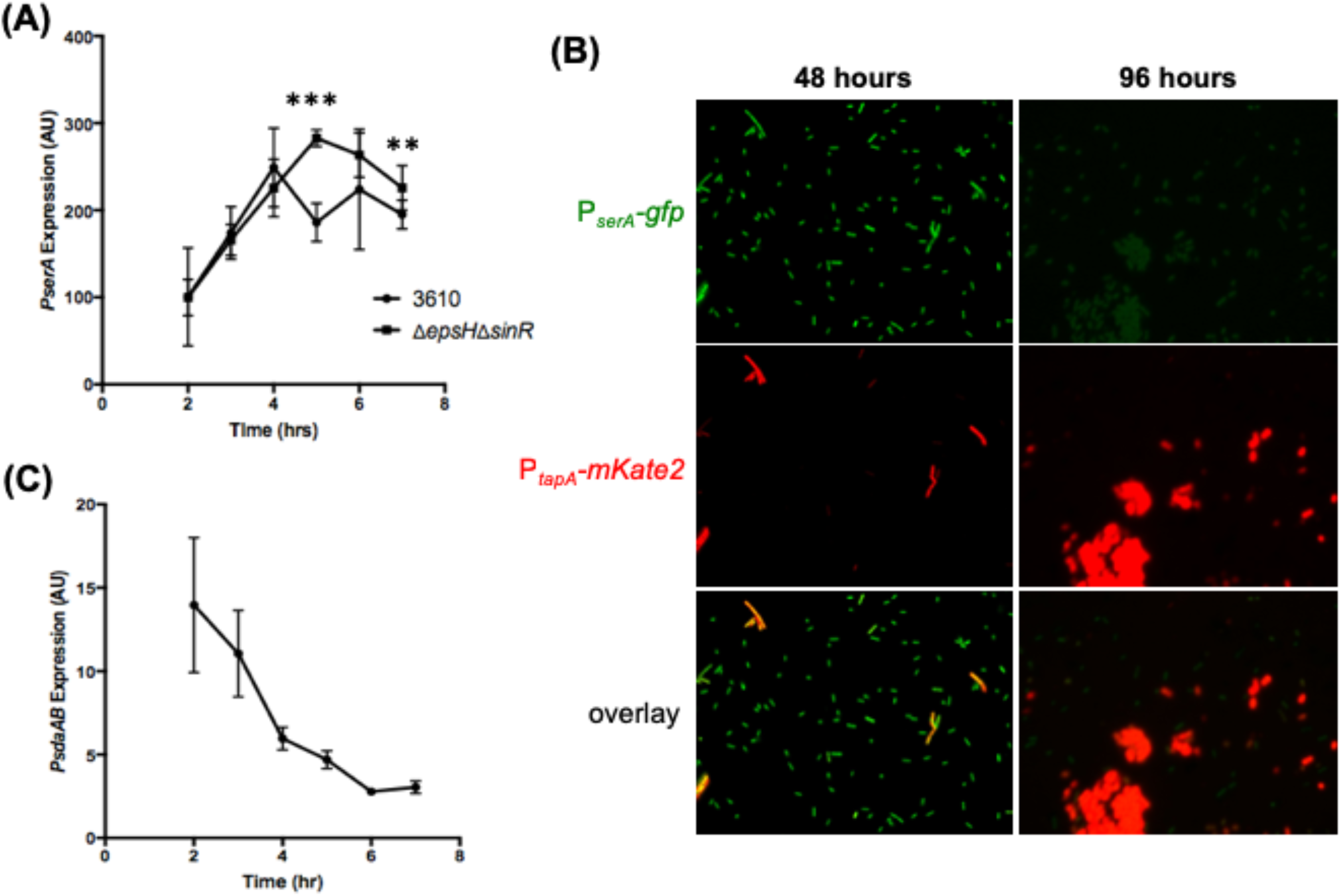
Expression of serine pathway in relation to biofilm gene expression. **(A)** The deletion of *sinR* locks cells into a “biofilm on” state (50), and to prevent clumping, *epsH*, a transferase gene involved in building the biofilm matrix (51) was also deleted. Beta-galactosidase assay quantification of *PserA* expression in wildtype cells (JG106) (circle) compared to the biofilm double mutant Δ*sinR*Δ*epsH* (ABR167) (square) under shaking conditions in MSgg medium. Wild type had peak expression at 4hrs, while the mutant peaked at 5hrs indicating a slight delay in expression decline. There is no significant difference in expression except at T5 and T7 (**=p<0.01, ***= p<0.001). The lack of significant difference (p>0.05) in other timepoints further suggests that expression of *serA* is regulated independent of biofilm formation. **(B)** Dual reporter assay of P*_serA_-GFP* (green) and P*_tapA_-mkate2* (red) in wildtype cells (JG157) during growth in MSgg medium. P*_serA_* expression was constitutively on at 48 hours but decreased significantly when cells entered mature biofilm stage (96 hours). The decrease was accompanied by the continued turn on of matrix gene expression (P*_tapA_-mkate2*). **(C)** Beta-galactosidase assay quantification of *PsdaA* expression in wildtype cells (ABR176) under shaking conditions in MSgg medium. Low overall expression and decreasing expression as entering biofilm stage.

Because the timing of decrease in *serA* expression coincided with the induction of biofilm genes, we wanted to further investigate a possible correlation between decreasing *serA* expression and biofilm gene induction. To do this, we constructed a dual fluorescent reporter strain. We fused the regulatory region of *serA* to the *gfp* gene, and the regulatory region of *tapA* to the *mKate2* gene, which encodes a red fluorescent protein, respectively. We then tracked the expression of the two reporter fusions in the engineered cell population over time during biofilm development in MSgg. As shown in Fig. 4B, most cells were expressing *serA* at 48 hours while only a small proportion of cells were expressing *tapA* (Fig. 4B). The bimodal pattern of *tapA* expression was reported previously by several published studies (43, 48, 49). After 96 hours, cells, many of which were in the form of bundles and clumps due to matrix production, were still strongly expressing *tapA*, while *serA* expression was largely deceased (Fig. 4B). Our results seemed to support the correlation between decrease in *serA* expression and the induction of biofilm genes although it was not yet clear if decreasing *serA* expression proceeded the biofilm gene induction.

We also wanted to investigate if the biofilm pathway impacts serine homeostasis through a putative feedback mechanism. To do so, we “locked” the cells into the “biofilm ON” state by deleting the *sinR* repressor gene. This allows matrix genes to be constitutively expressed for biofilm formation and results in hyper robust biofilms (50). The *epsH* gene, which encodes a sugar transferase for EPS biosynthesis, was also deleted to prevent cell clumping due to excess matrix production and facilitate sample collection during bioassays (51). Interestingly, there was no difference in *serA* expression up until the transition point (hour 4, Fig 4A). *serA* expression was decreased in the wild type cells while the “biofilm ON” cells did not decrease *serA* expression until an hour later (Fig. 4A). This result suggested that being in the “biofilm ON” state did not impact serine biosynthetic gene expression and that less likely decreasing serine levels was due to feedback regulation from the biofilm pathway.

Lastly, we wanted to examine the expression of the serine deaminase genes (*sdaAB-sdaAA*). If serine levels are being depleted, it may be as well because the serine deaminase is activated to convert existing serine back to pyruvate for metabolic purposes in addition to decreased serine biosynthesis. We tested this by constructing a *lacZ* fusion to the promoter of the *sdaA* operon (*sdaAB-sdaAA*), P*_sdaAB_-lacZ*. Cells were grown in MSgg medium and expression of the reporter was measured overtime. We found that overall *sdaA* expression was very low and declining over time (Fig. 4C). This result suggests that if the activity of the serine deaminase in the shunt pathway is upregulated it most likely occurs at the protein level, instead of gene expression. All together, our data suggest that during the transition from exponential phase to stationary phase growth, decreasing serine biosynthesis could be a very important factor influencing serine homeostasis and intracellular serine levels. The impact from the serine shunt pathway mediated is also substantial based on the biofilm phenotype and microbiological assays, while any putative regulation on the deaminase activity is still not clear.

### CcpA Activates *serA* Expression

Following the observation that *serA* expression was decreased upon entry into stationary phase, we decided to characterize potential regulators of *serA*. The regulation of *serA* is well-studied in *E. coli*, where serine biosynthesis is under the regulation of catabolite control, with *serA* being activated by both CRP and Lrp (52), indicating that its production increases when cells run out of preferred carbon sources and begin to rely on secondary carbon sources. In *B. subtilis*, CcpA, the master regulator of catabolite control, is a functional counterpart of CRP in *E. coli*. While commonly thought of as a transcriptional repressor (53, 54), there is evidence of CcpA also serving as an activator of genes in the pathways involved in carbon utilization, such as the acetate metabolism (55), or branched chain amino acid biosynthesis (leucine, isoleucine, and valine) (45, 56). We next examined levels of *serA* expression in a strain deleted for ccpA. We found that deletion of *ccpA* led to lowered levels of *serA* expression when compared to the wild type strain (Fig. 5A). Lowered expression of *serA* in the *ccpA* mutant was also confirmed using qRT-PCR (Fig. 5B). These results suggest that CcpA positively regulates *serA* expression. Interestingly, a previous study has shown that decreased CcpA activity led to biofilm induction in *B. subtilis* (27). The decreased *serA* expression and therefore serine levels upon decreased CcpA activity could be at least part of the mechanism by which the biofilm induction occurs.

**Figure 5:**
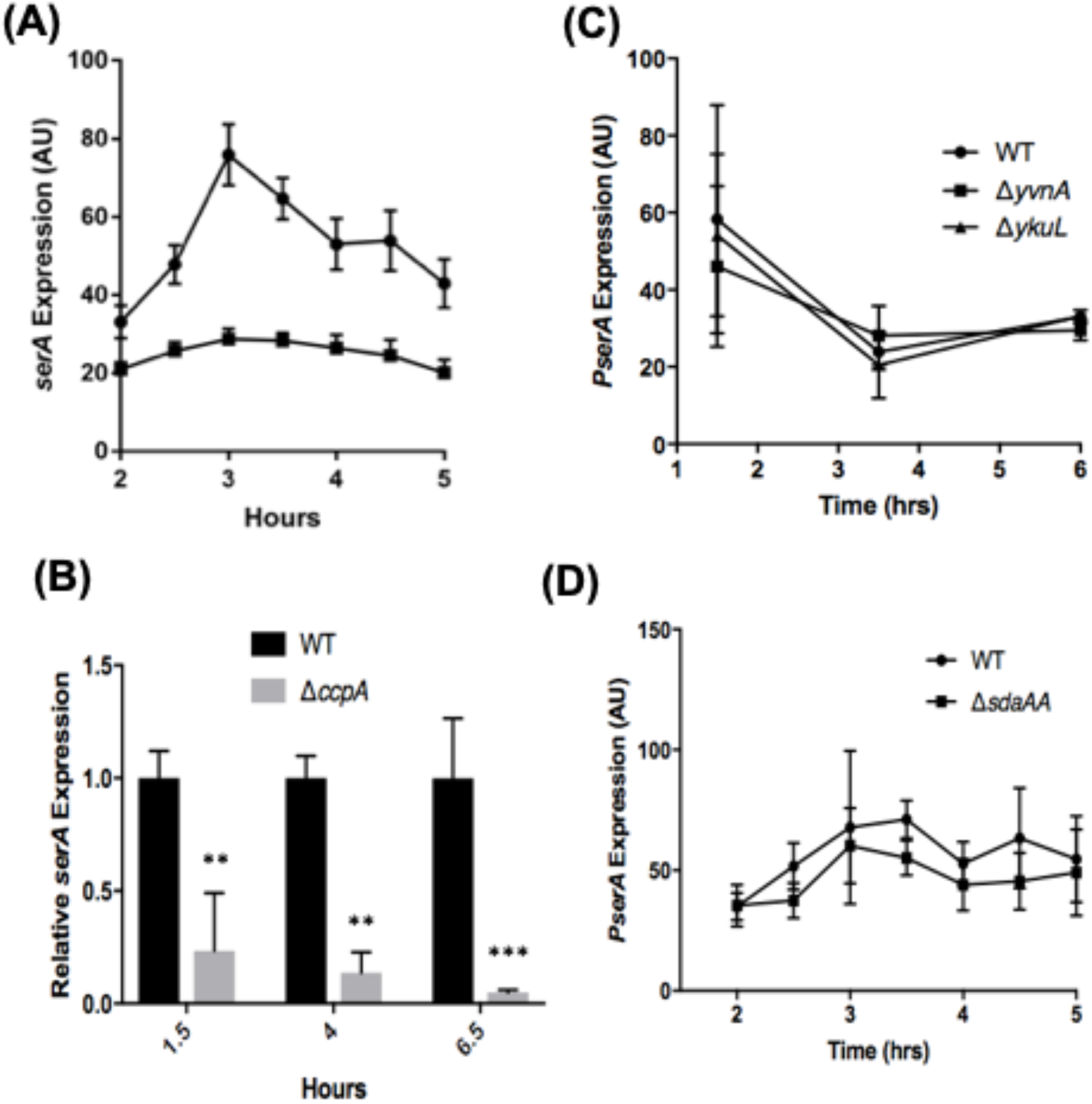
P*serA* transcription is regulated by CcpA but not intracellular serine levels. **(A)** Expression of *serA* in wildtype cells (circles) initially increased during exponential phase and then began to decrease at approximately OD_600_=1.00; arrow indicates OD_600_=1.00). Deletion of *ccpA (*squares) led to constitutively low expression of *serA*. **(B)** Confirmation of the β-galactosidase assay with qRT-PCR. At all time-points tested, *serA* mRNA levels were significantly lower than in the *DccpA* mutant (**=p≤0.01; ***=p≤0.001). **(C)** Two putative regulators, YvnA and YkuL, do not seem to regulate *serA* transcription. Genes *yvnA* (squares), and *ykuL* (triangles) were tested as potential regulators of *serA*. Deletion of neither gene showed a difference in expression relative to wildtype. **(D)** Deletion of the *sdaAA* gene did not significantly impact *serA* expression in cells either (p>0.05).

Since we could not find a direct binding site of CcpA in the *serA* promoter (data not shown) we speculated that CcpA regulates *serA* likely through repression of a repressor. In an effort to further elucidate the mechanism of *serA* regulation by CcpA, we examined two genes in the characterized CcpA regulon both encoding regulatory proteins of unknown function. Importantly, these two genes were shown to be upregulated under biofilm conditions (57–59). The first candidate gene, *ykuL*, encodes a putative CBS-domain containing protein. *ykuL* is both repressed by CcpA and activated by Spo0A (59, 60). The second candidate, *yvnA*, encodes a transcriptional repressor. *yvnA* is repressed by both CcpA and AbrB (57, 58). However as shown in Fig. 5C, deletion of neither gene had an impact on *serA* expression. Thus, how CcpA activates *serA* is still unclear. Lastly, deletion of *sdaAA* also did not significantly affect expression of *serA*(Fig. 5D), supporting the idea that serine levels do not affect *serA* transcription via any putative feedback mechanism.

### tRNA^ser^ Isoacceptors Are Expressed at Lower Levels in Stationary Phase Relative to Exponential Phase

In our previous study, we showed that serine starvation led to ribosomes preferentially pausing on the four UCN (N being U, A, C, G) serine codons, but not the AGC or AGU serine codon (16). The logical connection between intracellular serine levels and ribosome translation efficiency is the serine tRNAs that add the amino acid to the growing polypeptide chain within the ribosome. *B. subtilis* encodes a single seryl-tRNA-synthetase to carry out the charging of all five serine tRNA isoacceptors (33). The five isoacceptors are clustered into two subgroups, with one group (*trnB-ser2* and *trnS-ser*) recognizing AGC and AGU codons and the other (*trnB-ser1, trnD-ser*, and *trnSL-ser1*) recognizing the four UCN codons (Table 3). Given that there is only one synthetase for serine tRNA charging, we reasoned that differential activity/expression of tRNA isoacceptors could play a role in mediating the codon-specific ribosome pausing previously observed (16). By using qRT-PCR, we examined levels of the five tRNA^ser^ isoacceptors in both exponential phase and stationary phase cells. We observed that the levels of all five isoacceptors were decreased in stationary phase when compared to those in exponential phase (decrease ranged from ~37% to ~3%, stationary vs exponential), relative to the internal control of 16S rRNA (Fig. 6). Reduced levels of tRNA^ser^ isoacceptors in stationary phase *B. subtilis* cultures were indeed noted decades ago (At the time, it was speculated that there were three tRNA^ser^ isoacceptors base on size-fractionation technique)(61). Interestingly, our results also showed that isoacceptors recognizing the UCN serine codons (*trnB-ser1, trnD-ser*, and *trnSL-ser1*) were expressed at far lower levels than those recognizing AGY codons (*trnB-ser2* and *trnS-ser*) in stationary phase (Fig. 6). This significantly differentiated decrease in the levels of the tRNA^ser^ isoacceptors recognizing the four UCU serine codons and of those recognizing the AGC and AGU serine codons could contribute to the preferential pausing of ribosomes on the UCN serine codons under serine starvation that we observed previously.

**Figure 6:**
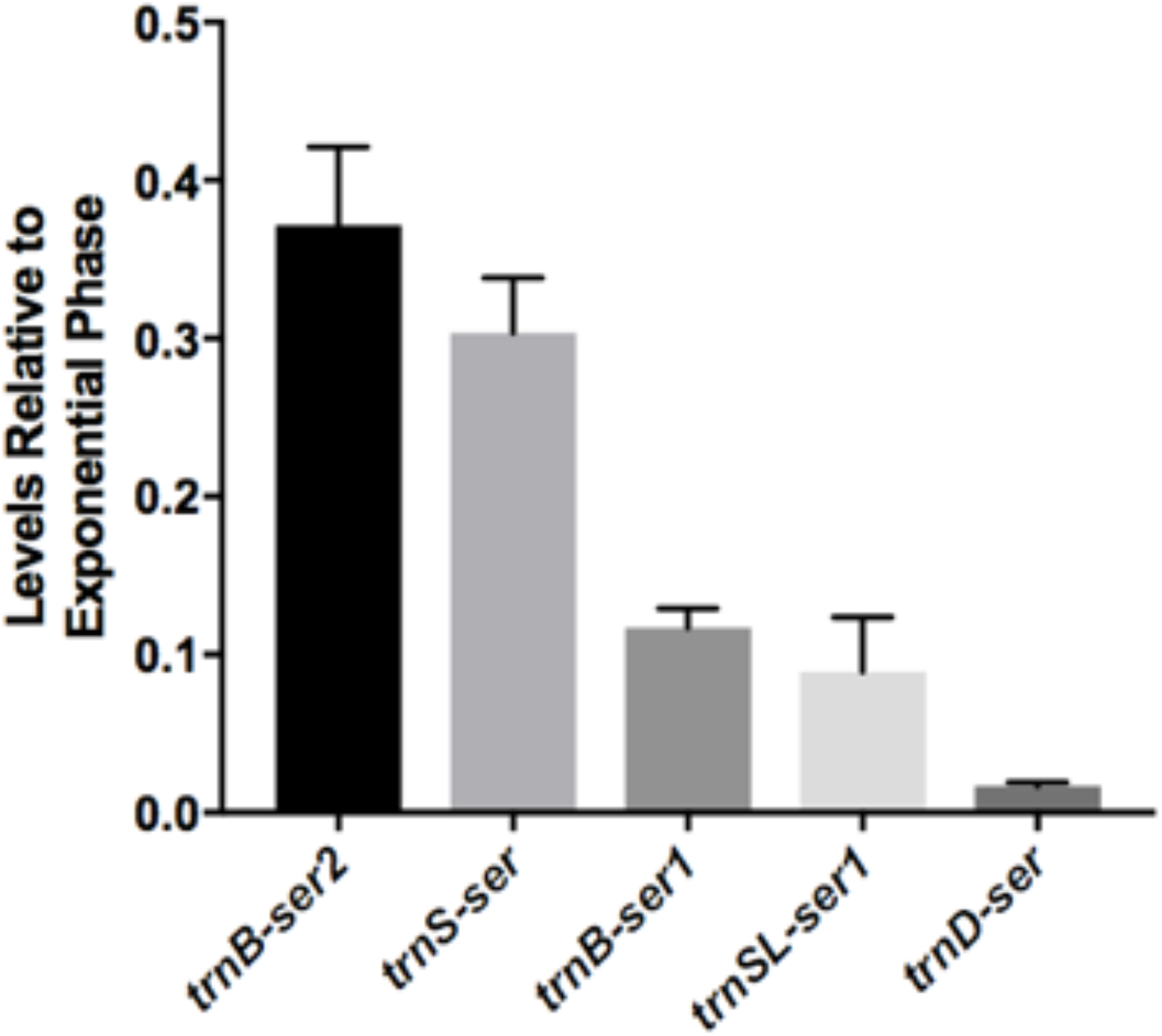
tRNA^ser^ isoacceptor transcription levels are decreased in stationary phase relative to exponential phase. tRNA^ser^ isoaccepter gene expression was quantified by RT-qPCR on cells grown to exponential and stationary phase in MSgg medium. There was a significant reduction in expression levels all five tRNA^ser^ isoacceptor genes in stationary phase relative to exponential phase. The three that recognize UCN codons *(trnB-ser1, trnSL-ser1, trnD-ser)* were found to have expression levels lower than those that recognize AGY codons. Error bars represent 95% confidence interval.

**Table 3:**
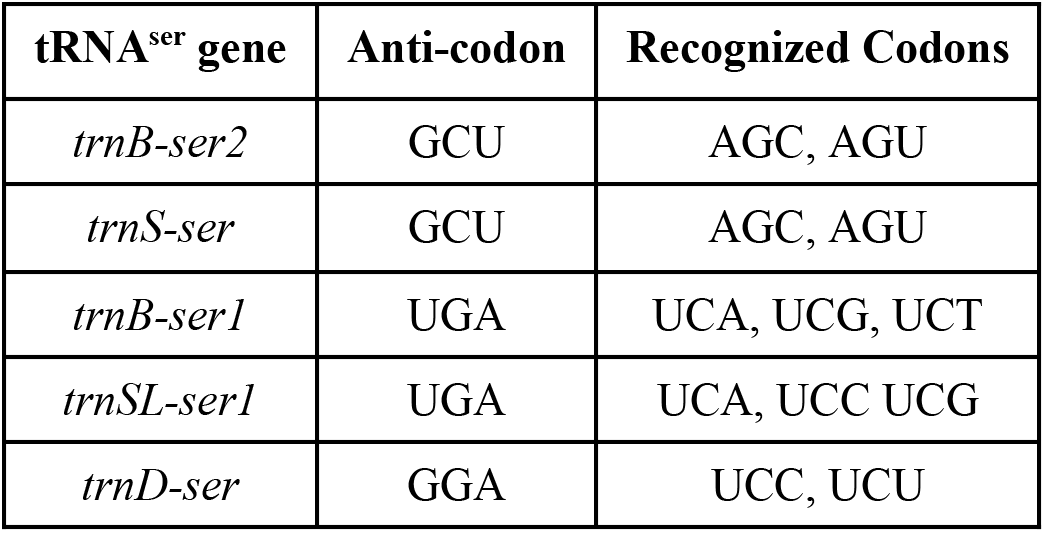
List of tRNA^ser^ isoacceptors in *B. subtilis* and the codons recognized by each isoacceptor.

| tRNA<sup>ser</sup> gene | Anti-codon | Recognized Codons |
| --- | --- | --- |
| *trnB-ser2* | GCU | AGC, AGU |
| *trnS-ser* | GCU | AGC, AGU |
| *trnB-ser1* | UGA | UCA, UCG, UCT |
| *trnSL-ser1* | UGA | UCA, UCC UCG |
| *trnD-ser* | GGA | UCC, UCU |

Following the above, we further argued that if ribosome pausing on selective serine codons is due to differentially lowed production of individual tRNA^ser^ isoacceptors upon serine starvation, it is then possible to be overcome by overexpressing corresponding tRNA^ser^ isoacceptors. As a prior example, it was shown that in *E. coli*, amino acid starvation led to differential charging of leucine tRNA isoacceptor*s*, which can be overcome by overexpressing the leucine tRNAs that are charged at lower levels during starvation (62). To test if our argument was valid, we used a genetic approach and overexpressed each of the five serine tRNA genes from an alternative chromosomal locus in the wild type strain. We then tested the affect of serine tRNA gene overexpression on matrix gene expression and on the timing of biofilm induction. However, our results showed that overexpression of the individual tRNAs did not impact matrix gene expression (Fig. S1A) nor the timing of biofilm induction (data not shown). It is possible however, that the induction was not strong enough to be physiologically relevant, as tRNAs are in general expressed at high levels in the cell and are quite stable (63, 64). Note that in the *E. coli* study, leucine tRNA genes were overexpressed using a high-copy-number plasmid system (65).

We also investigated the possibility that the wild-type strain was not sensitive enough, and thus used strains with synonymous substitutions in selected serine codons in the *sinR* gene to repeat serine tRNA gene overexpression. It was shown previously that synonymously mutating the AGC and AGT serine codons in the *sinR* gene to any of the four TCN codons dramatically increased the impact of serine starvation on *sinR* translation and increased expression of the matrix operon (16). However, overexpression of individual tRNA^ser^ genes in two engineered strains each carrying a different type of synonymous mutations in *sinR* (AGY to TCC in Fig. S1B and AGY to TCT in Fig. S1C, respectively) did not significantly affect matrix gene expression either (Fig. S1B-C).

## DISCUSSION

We previously showed that serine depletion triggered biofilm induction in *B. subtilis* due to global ribosome pausing on selective serine codons during translation and specifically a decrease in translation of the master biofilm repressor gene *sinR* (16). What remained unaddressed was the regulation driving the decrease in intracellular serine levels at the entry of stationary phase and the mechanism of serine starvation-triggered preferential ribosome pausing on the UCN serine codons. In this study, we investigated both how serine depletion occurs in early stationary phase (biofilm entry) and how changes in serine tRNA levels may be playing a role in causing ribosomes to stall selectively on UCN serine codons. By using a novel microbiological assay, we showed that levels of serine decrease when cells enter stationary phase. Our results suggest that serine levels in the cell act as a signal for biofilm formation. Our result also aligns with previous evidence that serine is one of the first amino acids to be exhausted in *E. coli* (34–36). While using supernatant is an indirect measure of intracellular levels, the finding that higher levels of auxotroph growth were supported when grown in supernatant from cells overexpressing *serA* or lacking *sdaAA*, supports there being a correlation between intracellular and extracellular serine levels (Fig. 3).

Serine depletion upon entry into stationary phase is likely due to a decrease in serine biosynthesis and the shunt pathway that converts serine to pyruvate. We also showed that the key serine biosynthesis gene *serA* is regulated by the catabolite repressor, CcpA, linking the regulation of serine homeostasis to cell global metabolic status. A previous study presented evidence suggesting that carbon metabolism and *serA* expression are linked, as levels of *serA* transcription decreased 22-fold following a minute of glucose starvation in M9 media (66), but the mechanism was unclear. Here we show that *serA* is subject to catabolite regulation, being activated by CcpA. This regulation is most likely indirect, due to the lack of a direct binding site of CcpA in the promoter region of *serA*. While most amino acid biosynthesis genes are upregulated under nutrient starvation, our data indicate that *serA* actually begins to decrease in expression upon entry into stationary phase, with peak expression happening at approximately the same time as *B. subtilis* is beginning to run out of its preferred carbon sources and it switches to secondary carbon sources, known as the diauxic shift (67).

A link between glucose metabolism and biofilm formation has been previously identified. Glucose represses biofilm formation in a CcpA-dependent manner (27). Our work fits into this model, adding a connection between a decrease in CcpA activity and an increase in biofilm formation. A lack of glucose would decrease CcpA activity, leading to a decrease in *serA* expression, which in turn, decreases serine levels. This drop in serine causes a decrease in SinR protein levels due to ribosomes pausing on the numerous TCN codons in *sinR* (16). Decreases in free SinR derepresses matrix gene expression, leading to biofilm formation. Regulating biofilm formation in this manner allows *B. subtilis* to monitor its environment as well as its intracellular metabolic state.

Finally, we showed that all five tRNA^ser^ isoacceptors were accumulated at lower levels in stationary phase relative to exponential phase. More interestingly, the decrease differs significantly for different isoacceptors in that the three isoacceptors recognizing the UCN serine codons decreased more substantially than the two isoacceptors recognizing the AGC and AGU serine codons. In this study, we primarily focused on the differential expression by using qRT-PCR. One can argue that shutting off transcription of tRNA genes would not be fast enough to reduce tRNA levels due to the stability of the tRNA in cells (64). One mechanism that may regulate this stability is the numerous modifications of tRNAs (reviewed in (68)). Modifications of tRNAs have been well studied in both eukaryotic cells and model bacteria such as *E. coli*, but less is known in *B. subtilis* (reviewed in (68–70)). Only two tRNA modification enzymes have been studied, including two homologs of *E. coli* MiaB, YmcB and YqeV (71). In *E. coli*, MiaB is involved in methylthiolation of selected tRNAs (72). In *B. subtilis*, the gene that encodes YmcB is in an operon with *ymcA*, a regulatory gene known to be essential for biofilm formation (50, 73, 74). In contrast to *ymcA*, the *ymcB* gene is not important for biofilm formation (YC, unpublished). Deletion of the *ymcB* gene also did not affect tRNA^ser^ levels (Data not shown). YmcB is a methylthiotransferase (MTT), which modifies an already present tRNA modification (71). It is possible that the primary modification is critical for regulating the degradation of tRNAs upon nutrient starvation.

In addition to gene expression, the decrease in serine tRNA levels may also be due to lack of charging. Recent work in *E. coli* suggests that tRNAs become unstable during amino acid starvation (75). Using strains auxotrophic for specific amino acids, it was shown that upon induction of starvation, levels of tRNAs decreased. This was noted for the specific amino acid cells were starved for as well as other amino acids. It was speculated that a reduction in charging (due to amino acid starvation) leads to decreased translation, which in turn triggers a degradation of excess tRNA, regardless of its acylation state (75). Our data fit into this hypothesis, as the decrease in serine tRNA expression levels coincides with a decrease in serine levels (Figs. 2B, 4A, and 5A).

## ACKNOWLEDGEMENTS

We thank the members of the Chai and Godoy labs for helpful discussions. We thank Leticia Lima Angelini, Christina Potter, Rachel Son, and Philip Wasson for technical assistance. Funding for this work was provided by Northeastern University and a grant from the National Science Foundation to Y. Chai (MCB1651732).

## AUTHOR CONTRIBUTIONS

JG, AR, KG, VG, and YC designed the experiments. JG, AR, and GD performed the experiments. TT provided substantial technical help. JG, AR, and YC wrote the manuscript.

## References

1. Hall-Stoodley L, Costerton JW, Stoodley P. 2004. Bacterial biofilms: from the natural environment to infectious diseases. Nat Rev Microbiol 2:95–108.

2. O’Toole G, Kaplan HB, Kolter R. 2000. Biofilm formation as microbial development. Annu Rev Microbiol 54:49–79.

3. Stoodley P, Sauer K, Davies DG, Costerton JW. 2002. Biofilms as complex differentiated communities. Annu Rev Microbiol 56:187–209.

4. Hall-Stoodley L, Stoodley P. 2009. Evolving concepts in biofilm infections. Cell Microbiol 11:1034–1043.

5. Branda SS, Gonzalez-Pastor JE, Ben-Yehuda S, Losick R, Kolter R. 2001. Fruiting body formation by Bacillus subtilis. Proc Natl Acad Sci U S A 98:11621–11626.

6. Vlamakis H, Chai Y, Beauregard P, Losick R, Kolter R. 2013. Sticking together: building a biofilm the Bacillus subtilis way. Nat Rev Microbiol 11:157–168.

7. Chen Y, Cao S, Chai Y, Clardy J, Kolter R, Guo JH, Losick R. 2012. A Bacillus subtilis sensor kinase involved in triggering biofilm formation on the roots of tomato plants. Mol Microbiol 85:418–430.

8. Chen Y, Yan F, Chai Y, Liu H, Kolter R, Losick R, Guo JH. 2013. Biocontrol of tomato wilt disease by Bacillus subtilis isolates from natural environments depends on conserved genes mediating biofilm formation. Environ Microbiol 15:848–864.

9. Beauregard PB, Chai Y, Vlamakis H, Losick R, Kolter R. 2013. Bacillus subtilis biofilm induction by plant polysaccharides. Proc Natl Acad Sci U S A 110:E1621–1630.

10. McLoon AL, Kolodkin-Gal I, Rubinstein SM, Kolter R, Losick R. 2011. Spatial Regulation of Histidine Kinases Governing Biofilm Formation in *Bacillus subtilis*. J. Bacteriology 193:679–685.

11. Hamon MA, Lazazzera BA. 2001. The sporulation transcription factor Spo0A is required for biofilm development in Bacillus subtilis. Mol Microbiol 42:1199–1209.

12. Burbulys D, Trach KA, Hoch JA. 1991. Initiation of sporulation in B. subtilis is controlled by a multicomponent phosphorelay. Cell 64:545–552.

13. Chai Y, Norman T, Kolter R, Losick R. 2011. Evidence that metabolism and chromosome copy number control mutually exclusive cell fates in Bacillus subtilis. EMBO J 30:1402–1413.

14. Bai U, Mandic-Mulec I, Smith I. 1993. SinI modulates the activity of SinR, a developmental switch protein of Bacillus subtilis, by protein-protein interaction. Genes Dev 7:139–148.

15. Newman JA, Rodrigues C, Lewis RJ. 2013. Molecular Basis of the Activity of SinR Protein, the Master Regulator of Biofilm Formation in Bacillus subtilis. Journal of Biological Chemistry 288:10766–10778.

16. Subramaniam AR, Deloughery A, Bradshaw N, Chen Y, O’Shea E, Losick R, Chai Y. 2013. A serine sensor for multicellularity in a bacterium. Elife 2:e01501.

17. Townsley L, Yannarell SM, Huynh TN, Woodward JJ, Shank EA. 2018. Cyclic di-AMP Acts as an Extracellular Signal That Impacts Bacillus subtilis Biofilm Formation and Plant Attachment. MBio 9.

18. Chen Y, Chai Y, Guo JH, Losick R. 2012. Evidence for cyclic Di-GMP-mediated signaling in Bacillus subtilis. J Bacteriol 194:5080–5090.

19. Stauffer GV. 2004. Regulation of Serine, Glycine, and One-Carbon Biosynthesis. EcoSal Plus 1.

20. Saski R, Pizer LI. 1975. Regulatory properties of purified 3-phosphoglycerate dehydrogenase from Bacillus subtilis. Eur J Biochem 51:415–427.

21. Rex JH, Aronson BD, Somerville RL. 1991. The tdh and serA operons of Escherichia coli: mutational analysis of the regulatory elements of leucine-responsive genes. 173:5944–5953.

22. de Crombrugghe B, Busby S, Buc H. 1984. Cyclic AMP receptor protein: role in transcription activation. Science 224:831–838.

23. Taniguchi T, O’Neill M, de Crombrugghe B. 1979. Interaction site of Escherichia coli cyclic AMP receptor protein on DNA of galactose operon promoters. Proc Natl Acad Sci U S A 76:5090–5094.

24. Newman EB, D’Ari R, Lin RT. 1992. The leucine-Lrp regulon in E. coli: a global response in search of a raison d’etre. Cell 68:617–619.

25. Henkin TM, Grundy FJ, Nicholson WL, Chambliss GH. 1991. Catabolite repression of alpha-amylase gene expression in Bacillus subtilis involves a trans-acting gene product homologous to the Escherichia coli lacl and galR repressors. Mol Microbiol 5:575–584.

26. Gorke B, Stulke J. 2008. Carbon catabolite repression in bacteria: many ways to make the most out of nutrients. Nat Rev Microbiol 6:613–624.

27. Stanley NR, Britton RA, Grossman AD, Lazazzera BA. 2003. Identification of catabolite repression as a physiological regulator of biofilm formation by Bacillus subtilis by use of DNA microarrays. J Bacteriol 185:1951–1957.

28. Chen Y, Gozzi K, Yan F, Chai Y. 2015. Acetic acid acts as a volatile signal to stimulate bacterial biofilm formation. mBio 6:e00392-00315.

29. Brinsmade SR, Alexander EL, Livny J, Stettner AI, Segre D, Rhee KY, Sonenshein AL. 2014. Hierarchical expression of genes controlled by the *Bacillus subtilis* global regulatory protein CodY. P Natl Acad Sci USA 111:8227–8232.

30. Su HS, Lang BF, Newman EB. 1989. L-serine degradation in Escherichia coli K-12: cloning and sequencing of the sdaA gene. J Bacteriol 171:5095–5102.

31. Tobey KL, Grant GA. 1986. The nucleotide sequence of the serA gene of Escherichia coli and the amino acid sequence of the encoded protein, D-3-phosphoglycerate dehydrogenase. J Biol Chem 261:12179–12183.

32. Zhang X, Newman E. 2008. Deficiency in l-serine deaminase results in abnormal growth and cell division of Escherichia coli K-12. Mol Microbiol 69:870–881.

33. Zhu B, Stulke J. 2018. SubtiWiki in 2018: from genes and proteins to functional network annotation of the model organism Bacillus subtilis. Nucleic Acids Res 46:D743–D748.

34. Liebs P, Riedel K, Graba JP, Schrapel D, Tischler U. 1988. Formation of some extracellular enzymes during the exponential growth of Bacillus subtilis. Folia Microbiol (Praha) 33:88–95.

35. Sezonov G, Joseleau-Petit D, D’Ari R. 2007. Escherichia coli physiology in Luria-Bertani broth. J Bacteriol 189:8746–8749.

36. Pruss BM, Nelms JM, Park C, Wolfe AJ. 1994. Mutations in NADH:ubiquinone oxidoreductase of Escherichia coli affect growth on mixed amino acids. J Bacteriol 176:2143–2150.

37. Shemesh M, Chai Y. 2013. A combination of glycerol and manganese promotes biofilm formation in Bacillus subtilis via histidine kinase KinD signaling. J Bacteriol 195:2747–2754.

38. Sambrook J. 2001. Molecular Cloning. A Laboratory Manual. Cold Spring Harbor Laboratory Press Cold Spring Harbor, NY, USA.

39. Gryczan TJ, Contente S, Dubnau D. 1978. Characterization of Staphylococcus aureus plasmids introduced by transformation into Bacillus subtilis. J Bacteriol 134:318–329.

40. Yasbin RE, Young FE. 1974. Transduction in Bacillus subtilis by bacteriophage SPP1. J Virol 14:1343–1348.

41. Chai Y, Kolter R, Losick R. 2009. A widely conserved gene cluster required for lactate utilization in Bacillus subtilis and its involvement in biofilm formation. J Bacteriol 191:2423–2430.

42. Antoniewski C, Savelli B, Stragier P. 1990. The spoIIJ gene, which regulates early developmental steps in Bacillus subtilis, belongs to a class of environmentally responsive genes. J Bacteriol 172:86–93.

43. Chai Y, Chu F, Kolter R, Losick R. 2008. Bistability and biofilm formation in Bacillus subtilis. Mol Microbiol 67:254–263.

44. Gozzi K, Ching C, Paruthiyil S, Zhao Y, Godoy-Carter V, Chai Y. 2017. Bacillus subtilis utilizes the DNA damage response to manage multicellular development. NPJ Biofilms Microbiomes 3:8.

45. Molle V, Nakaura Y, Shivers RP, Yamaguchi H, Losick R, Fujita Y, Sonenshein AL. 2003. Additional targets of the Bacillus subtilis global regulator CodY identified by chromatin immunoprecipitation and genome-wide transcript analysis. J Bacteriol 185:1911–1922.

46. Chu F, Kearns DB, McLoon A, Chai Y, Kolter R, Losick R. 2008. A novel regulatory protein governing biofilm formation in Bacillus subtilis. Mol Microbiol 68:1117–1127.

47. DeLoughery A, Lalanne J-B, Losick R, Li G-W. 2018. Maturation of polycistronic mRNAs by the endoribonuclease RNase Y and its associated Y-complex in Bacillus subtilis. PNAS 115:E5585–E5594.

48. Vlamakis H, Aguilar C, Losick R, Kolter R. 2008. Control of cell fate by the formation of an architecturally complex bacterial community. Genes Dev 22:945–953.

49. Lopez D, Fischbach MA, Chu F, Losick R, Kolter R. 2009. Structurally diverse natural products that cause potassium leakage trigger multicellularity in Bacillus subtilis. Proc Natl Acad Sci U S A 106:280–285.

50. Kearns DB, Chu F, Branda SS, Kolter R, Losick R. 2005. A master regulator for biofilm formation by Bacillus subtilis. Mol Microbiol 55:739–749.

51. Marvasi M, Visscher PT, Casillas Martinez L. 2010. Exopolymeric substances (EPS) from Bacillus subtilis: polymers and genes encoding their synthesis. FEMS Microbiol Lett 313:1–9.

52. Yang L, RT Lin, EB Newman. 2002. Structure of the Lrp-regulated serA promoter of Escherichia coli K-12. . Molecular Microbiology 43:323–333.

53. Tuan LR, R. D’Ari, and E.B. Newman. 1990. The leucine regulon of Escherichia coli K-12: a mutation in rblA alters expression of L-leucine-dependent metabolic operons. . Journal of Bacteriology 172.

54. Hueck CJ, and W. Hillen. 1995. Catabolite repression in Bacillus subtilis: a global regulatory mechanism for the gram-positive bacteria? Molecular Microbiology 15:395–401.

55. Grundy FJ, Waters, D.A, Allen S.H., Henkin, T.M. 1993. Regulation of the Bacillus subtilis acetate kinase gene by CcpA. . Journal of Bacteriology 175:7348–7355.

56. Wunsche A, Hammer E, Bartholomae M, Volker U, Burkovski A, Seidel G, Hillen W. 2012. CcpA forms complexes with CodY and RpoA in Bacillus subtilis. FEBS J 279:2201–2214.

57. Chumsakul O, Takahashi H, Oshima T, Hishimoto T, Kanaya S, Ogasawara N, Ishikawa S. 2011. Genome-wide binding profiles of the Bacillus subtilis transition state regulator AbrB and its homolog Abh reveals their interactive role in transcriptional regulation. Nucleic Acids Res 39:414–428.

58. Blencke HM, Homuth G, Ludwig H, Mader U, Hecker M, Stulke J. 2003. Transcriptional profiling of gene expression in response to glucose in Bacillus subtilis: regulation of the central metabolic pathways. Metab Eng 5:133–149.

59. Molle V, Fujita M, Jensen ST, Eichenberger P, Gonzalez-Pastor JE, Liu JS, Losick R. 2003. The Spo0A regulon of Bacillus subtilis. Mol Microbiol 50:1683–1701.

60. Blencke HM, Reif I, Commichau FM, Detsch C, Wacker I, Ludwig H, Stulke J. 2006. Regulation of citB expression in Bacillus subtilis: integration of multiple metabolic signals in the citrate pool and by the general nitrogen regulatory system. Arch Microbiol 185:136–146.

61. Doi RH, Kaneko I, Goehler B. 1966. Regulation of a serine transfer RNA of Bacillus subtilis under two growth conditions. Proc Natl Acad Sci U S A 56:1548–1551.

62. Sorensen MA, Elf J, Bouakaz E, Tenson T, Sanyal S, Bjork GR, Ehrenberg M. 2005. Over expression of a tRNA(Leu) isoacceptor changes charging pattern of leucine tRNAs and reveals new codon reading. J Mol Biol 354:16–24.

63. Waldron CaFL. 1975. Effect of growth rate on the amounts of ribosomal and transfer ribonucleic acids in yeast. Journal of Bacteriology 122:855–865.

64. Davis BD, Luger SM, Tai PC. 1986. Role of ribosome degradation in the death of starved Escherichia coli cells. J Bacteriol 166:439–445.

65. Subramaniam AR, Pan T, Cluzel P. 2013. Environmental perturbations lift the degeneracy of the genetic code to regulate protein levels in bacteria. Proc Natl Acad Sci U S A 110:2419–2424.

66. de Jong IG, Veening JW, Kuipers OP. 2012. Single cell analysis of gene expression patterns during carbon starvation in Bacillus subtilis reveals large phenotypic variation. Environ Microbiol 14:3110–3121.

67. Jacob F, Monod J. 1961. Genetic regulatory mechanisms in the synthesis of proteins. J Mol Biol 3:318–356.

68. Shepherd J, Ibba M. 2015. Bacterial transfer RNAs. FEMS Microbiol Rev 39:280–300.

69. Pan T. 2018. Modifications and functional genomics of human transfer RNA. Cell Res 28:395–404.

70. Maraia RJ, Arimbasseri AG. 2017. Factors That Shape Eukaryotic tRNAomes: Processing, Modification and Anticodon-Codon Use. Biomolecules 7.

71. Anton BP, Russell SP, Vertrees J, Kasif S, Raleigh EA, Limbach PA, Roberts RJ. 2010. Functional characterization of the YmcB and YqeV tRNA methylthiotransferases of Bacillus subtilis. Nucleic Acids Res 38:6195–6205.

72. Esberg B, Leung H-CE, Tsui H-CT, Björk GR, Winkler ME. 1999. Identification of the <em>miaB</em> Gene, Involved in Methylthiolation of Isopentenylated A37 Derivatives in the tRNA of <em>Salmonella typhimurium</em> and<em>Escherichia coli</em>. J. Bacteriology 181:7256–7265.

73. Branda SS, Gonzalez-Pastor JE, Dervyn E, Ehrlich SD, Losick R, Kolter R. 2004. Genes involved in formation of structured multicellular communities by Bacillus subtilis. J Bacteriol 186:3970–3979.

74. DeLoughery A, Dengler V, Chai Y, Losick R. 2016. Biofilm formation by Bacillus subtilis requires an endoribonuclease-containing multisubunit complex that controls mRNA levels for the matrix gene repressor SinR. Mol Microbiol 99:425–437.

75. Svenningsen SL, M. Kongstad, T.S. Stenum, A.J. Munoz-Gomez, M.A. Sorensen 2017. Transfer RNA is highly unstable during early amino acid starvation in Escherichia coli. Nucleic Acids Research 45:793–804.

76. Schroeder JW, Simmons LA. 2013. Complete Genome Sequence of Bacillus subtilis Strain PY79. Genome Announc 1.

77. Kunst F, Ogasawara N, Moszer I, Albertini AM, Alloni G, Azevedo V, et al. 1997. The complete genome sequence of the gram-positive bacterium Bacillus subtilis. Nature 390:249–256.

